# Endothelial SHP-1 regulates diabetes-induced abnormal collateral vessel formation and endothelial cell senescence

**DOI:** 10.1101/2024.04.26.591415

**Authors:** Alexandre Nadeau, Marike Ouellet, Raphaël Béland, Clément Mercier, Stéphanie Robillard, Farah Lizotte, Marc-Antoine Despatis, C. Florian Benzinger, Pedro Geraldes

## Abstract

**Objective:** Ischemia due to narrowing of the femoral artery and distal vessels is a major cause of peripheral arterial disease and morbidity affecting patients with diabetes. Diabetes-induced premature senescence of endothelial cells (EC) has been proposed as a mechanism leading to impaired ischemia-driven angiogenesis. Importantly, our previous work has shown that hyperglycemia reduced vascular endothelial growth factor (VEGF) activity in ischemic muscle of diabetic mice, which was associated with increased expression of the protein tyrosine phosphatase Src homology region 2 domain-containing phosphatase-1 (SHP-1). Here, we evaluate the impact of SHP-1 deletion on EC function and senescence.

**Approach and Results:** Ligation of the femoral artery was performed in nondiabetic (NDM) and 3 months diabetic (DM) mice with EC-specific deletion of SHP-1 and blood flow reperfusion was measured for 4 weeks. Blood flow reperfusion and limb function during voluntary wheel running were reduced by 43% and 82%, respectively in DM mice as compared to NDM mice. EC-specific deletion of SHP-1 in DM mice restored blood flow reperfusion by 60%, limb function by 86%, while capillary density was similar to NDM mice. Moreover, ablation of SHP-1 in EC prevented diabetes-induced expression of the senescence markers p53 and p21 and counteracted Nrf2 downregulation. In cultured EC, overexpression of dominant negative of SHP-1 prevented HG-induced inhibition of proliferation, migration, tubule formation and VEGFR2/Akt phosphorylation following VEGF stimulation. In addition, the expression of senescence markers and suppression of Nrf2 in EC exposed to HG levels were reversed by overexpression of dominant negative SHP-1.

**Conclusion:** SHP-1 in ECs is a central effector of diabetes-induced senescence that blocks VEGF action, and induces aberrant collateral vessel formation and blood flow reperfusion. Reduced SHP-1 expression counteracts these pathologic features suggesting the notion that it represents a promising therapeutic target.

**HIGHLIGHTS:**

- Endothelial specific deletion of SHP-1 (Scr homology 2-containing phosphatase-1) improves blood flow reperfusion, limb motricity and vessel density in the diabetic ischemic limb.
- Diabetes-induced SHP-1 protein expression inhibits VEGF proangiogenic actions and promotes endothelial senescence.
- Endothelial specific deletion of SHP-1 restores VEGF proangiogenic actions and prevents senescence in ischemic muscle and artery of diabetic mice and patients.

## INTRODUCTION

Peripheral artery disease (PAD) is a vascular complication associated with atherosclerotic plaque formation in the blood vessels of the lower extremities resulting in reduced blood flow and limb ischemia^1^. PAD is a major risk factor for lower extremity amputation, especially in patients with diabetes^2^. The disease normally presents with the appearance of painful muscle cramps in the calves during exercise that are usually relieved at rest, also known as intermittent claudication. The appearance of pain during the walking distance test is an indicator of PAD progression^3^. Unfortunately, about 50% of patients with PAD are asymptomatic and are therefore at higher risk of death, myocardial infarction, and heart attack^4^. The severity of PAD in patients with diabetes is strongly associated with duration and glycemic control^5^. During ischemia, collateral vessel development is insufficient to support the loss of blood flow through occluded arteries in patients with diabetic PAD^6^. Thus, therapeutic angiogenesis has been expected as a potential approach to treat patients with limb ischemia, but so far has shown limited clinical outcomes^7,8^. Several abnormalities in the angiogenic response to ischemia have been documented in the diabetic state involving complex interactions of multiple growth factors and vascular cells^9^.

The activation of growth factor receptors (such as VEGF and PDGF receptors) depends on the phosphorylation of their tyrosine residues, which are highly regulated by the balance between the level of protein tyrosine kinases and protein tyrosine phosphatases (PTPs)^10^. A dysregulation of the level of PTP expression can lead to dysfunction of cellular processes essential for blood vessel formation, including cell proliferation, migration, sprouting and differentiation^11^. The PTP superfamily is divided into two categories, phosphatases specifically recognizing phosphorylated tyrosine residues, known as classical PTPs, and dual-specificity phosphatases^12^. The Src homology region 2 domain-containing phosphatase-1 (SHP-1) is a cytoplasmic PTP containing two SH2 domains in the N-terminal of the protein as well as a phosphatase domain in the C-terminal that is activated by binding of the two SH2s to phosphorylated tyrosine residues causing a conformational change^13^. Unlike SHP-2, which can act as a positive or negative regulator, SHP-1 is only a negative regulator of several signaling pathways^14^. SHP-1 regulates the phosphorylation and activation of VEGFR-2 and PDGFR-β receptors^15^. Our group has reported that SHP-1 expression is elevated in the ischemic muscle of diabetic patients who underwent amputation following complications of PAD and in the ischemic muscle of diabetic mice following femoral artery ligation^16^. This increase in SHP-1 expression was associated with a decrease in PDGFR-β and Akt phosphorylation in the ischemic muscle of diabetic mice^17^. In endothelial cells exposed to high glucose levels, increased SHP-1 phosphatase activity was observed, which was associated with upstream activation of protein kinase C delta and angiotensin II receptor^18,19^. However, in addition to the regulation of growth factor signaling, SHP-1 could be involved in other processes such as premature cell cycle arrest and cellular senescence that contribute to endothelial dysfunction and poor angiogenic response to hypoxia in diabetes.

Senescence represents an irreversible arrest of the cell cycle, preventing cell proliferation. It can be triggered by various intrinsic or extrinsic stresses, including oxidative stress, DNA damage, telomere shortening, and genotoxic agents^20^. Senescence occurs mainly in aging and in the case of disease, it can have a positive or negative impact^21^. Its role was mainly investigated in cancer therapy development to prevent or limit the growth of cancer cells, but accumulation of cellular senescent cells can also decrease repair capacity of tissues in the elderly^22^. Vascular aging, caused by growing number of senescent cells, leads to endothelial structure and function alteration including angiogenic disorders, which increase the risk of vascular complications with age^23^. Protein expression associated with senescence such as p53 and p21 was observed in the ischemic muscle of mice on three days following femoral artery ligation and an increase in the amount of pro-senescent molecules was observed in the serum of patients with PAD^24^. Therefore, evidence suggests that both hyperglycemia and oxidative stress are important factors in the activation of senescence and may contribute to the endothelial dysfunction found in diabetic patients with PAD. However, the exact mechanism of diabetes-induced poor angiogenic function and accelerated senescence are not completely understood^25^. Given the central role of SHP-1 in PAD it is conceivable that it is also has senescence inducing functions.

Here we used diabetic and non-diabetic EC-specific knockout mice in combination with a loss-of-function cell culture paradigm to study the role of SHP-1 in high-glucose induced angiogenic dysfunction and cellular senescence. We observed that loss of SHP-1 protein or function counteracts senescence and diabetes-associated features in EC in vivo and in vitro identifying it as a central hub involved in disease pathogenesis.

## RESEARCH DESIGN AND METHODS

### Reagents and antibodies

All antibodies and reagent name, catalogue number and their suppliers are detailed in the Major Resources Table in the Supplemental Materials

### Human tissue study

After written consent under the approval of the Research Ethics Board of the CHUS (“Centre Hospitalier Universitaire de Sherbrooke”) [REB #2020-3107], the tibial artery and a biopsy of the gastrocnemius muscle of the ischemic limb were obtained from 9 patients (3 without and 6 with diabetes) that underwent surgery for lower limb amputation at the CHUS. Data were used to study SHP-1 levels in the human PAD. Patient characteristics are listed in Supplemental Table S1.

### Animals and experimental design

To investigate the role of SHP-1 specifically in endothelial cell, tamoxifen inducible mice C57BL/6-Cdh5-cre/ERT2 (no 13073 from Taconic) (specific to endothelial cells) were bred to SHP-1 flox mice (JAX stock no. 008336; B6.129P2-Ptpn6tm1Rsky/J). Diabetes was induced by intraperitoneal streptozotocin injection (90 mg/kg in 0.05 mol/L citrate buffer, pH 4.5; Sigma) on 2 consecutive days after 6-8 hours fasting at 7 weeks of age; control mice were injected with citrate buffer. Diabetes was confirmed the week after based on blood glucose levels measured with a Contour glucometer (Bayer). After 2 months of uncontrolled diabetes, 75mg/kg of tamoxifen (Toronto research chemicals) diluted in corn oil was injected intraperitoneally on 5 consecutive days to induce the endothelial-specific knockout of SHP-1. Femoral artery ligation was performed the following week. Throughout the study period, animals were provided with free access to water and standard rodent chow (Envigo Teklad). All experiments were conducted under the ethical protocol 2021-3170 in accordance with the Canadian Council of Animal Care, the University of Sherbrooke, and the NIH Guide for the Care and Use of Laboratory Animals.

### Assessment of endothelial specific SHP-1 deletion

Endothelial cells (EC) were extracted from the lungs of each group of mice. Lungs were sliced into 1-2 mm pieces and incubated at 37°C for 1 h in 0.2% collagenase type 1. The tissue was disrupted by pipetting and passed through a 40 μm cell strainer. Following centrifugation, the pellet was resuspended in 1 ml DMEM/0.1% BSA and incubated with CD31-conjugated Dynabeads for 30 min at 4°C. Beads (linked to EC) were washed with DMEM/0.1% BSA using a magnetic support. Smooth muscle cells (SMC) were isolated from the mouse aorta. Briefly, the aorta from each mouse was isolated and dissected to expose the inside surface for gentle removal of EC. The aorta was cut into small pieces and put face down in a sterile 24-well culture plate with DMEM 10% FBS. Following the appearance of SMC around the aorta, the tissue was removed to let SMC grow until confluence. EC and SMC were digested with proteinase K O/N at 55 °C in final concentration of 1X Modified Gitschier Buffer (Tris HCL pH 8.8 67 mM, Ammonium Sulfate 16 mM, MgCl2 6.5 mM) with 0.5% triton and 1% 2-beta-mercaptoethanol. Presence of *loxP* sites and transgene were validated by genotyping with specific primers listed in Supplemental Table S2. Recombination of *loxP* was validated using the same forward (F1) SHP-1 primer which target the 3’ loxP, and a second reverse primer (R2) which target is located after the second *loxP* sequence (see genotyping strategy in Supplemental Fig. S1).

*Hindlimb ischemia model* – To reproduce PAD and investigate revascularisation, ischemia of one lower limb was induced by ligation of the femoral artery in all four groups (NDM; DM; ec-SHP-1^+/+^, ec-SHP-1^-/-^) as we previously described^16^. Euthanasia was performed following laser Doppler perfusion imaging at day 28 by exsanguination via the heart left ventricle under deep anaesthesia (Isoflurane USP, inhalation at a concentration of 5%).

*Laser Doppler Perfusion Imaging and exercise willingness* – Hindlimb blood flow was measured using a laser Doppler perfusion imaging (PIMIII) system (Perimed Inc). Measurements were performed on shaved limb pre- and post-surgery, and additionally on days 7, 14, 21, and 28 post artery ligation, on anesthetized animals. All results are presented as a ratio of simultaneous measurement of the right (ligated) and left (non-ligated) limb to account for variables that can affect blood flow temporally. Exercise willingness was assessed in the last 5 days before euthanasia by allowing the mice voluntary access to activity wheel (Lafayette Instrument) in individual cages.

### Tissue preparation, histopathology and immunofluorescence

Right and left adductor muscles from nondiabetic mice; diabetic mice; ec-SHP-1^+/+^, ec-SHP-1^−/−^ mice were harvested for pathological analysis. The tissue sections (4-µm) sections were prepared and stained with hematoxylin & eosin or used for immunofluorescence as previously described^16^. The entire tissue sample of the muscle fiber structure of 6 or 7 mice per group were taken using Nikon TI-Eclipse microscope under identical conditions. Mean fiber diameter was obtained by measuring the diameter of 50 representative fibers of each animal with the Image J software^26^. Vessels with a diameter ranging from 10 to 30 μm were counted and considered as arterioles and endothelial cells without smooth muscle cells were counted as capillaries.

### Immunoblot analyses

Proteins from the adductors muscles or endothelial cells were lysed, extracted and treated as previously described^16,27^. Protein content quantification was performed using computer-assisted densitometry with ImageLab imaging software (Chemidoc, BioRad)

### Real-time PCR analyses

Real-time PCR was performed to obtain levels of mRNA expression of genes implicated in angiogenesis. Human vessels and mouse ischemic adductor muscle were incubated with 300 μl of Tri reagent and grinded using manual homogenization and RNA extraction was performed according to the manufacturers instructions. Reverse transcription and real-time PCR were performed as previously described^28^. PCR primers are listed in the Supplemental Table S3. GAPDH and B2M were used as housekeeping genes to normalize data.

### Cell culture and adenoviral infection

Bovine aortic endothelial cells (ECs) were isolated from freshly harvested aortas that were obtained from a local slaughterhouse as previously described^19,27^. Cells were cultured in DMEM 2.5% FBS, 1% penicillin-streptomycin and exposed to DMEM 0.1% FBS containing normal glucose (NG; 5.6 mmol/L + 19.4 mmol/L mannitol to adjust osmotic pressure) or high glucose (HG; 25 mmol/L) levels up to 48 h. To induce hypoxic conditions, cells were placed into an incubator set at 1% O2 for the last 16 hours and then stimulated with VEGF-a (10 ng/mL) for 5 minutes. Adenoviral vectors containing the dominant negative (Ad) form of SHP-1 (Ad-dnSHP-1) or GFP (green fluorescent protein; Ad-GFP) as control were used to infect BAEC as we have previously reported^15^. Cells from passages between 2 and 7 were used to study signaling and from passages between 9 to 14 to study senescence.

### Proliferation, migration and lumen formation assay

EC were infected with ad-GFP or Ad-dnSHP-1 and then exposed to NG or HG concentrations for 48 h with or without VEGF 25 ng/ml for 24 h (proliferation and migration) or 4 h (lumen formation) and on 1% O_2_ for the last 16 h. Proliferation, migration and lumen formation quantification were performed as previously described^27^.

### Β-galactosidase essay

Infected BAECs were seeded in 24-well plates at 30 000 cells per well for 4h, following exposure to NG or HG levels for 24 h and under hypoxic condition (1% O2) during the last 16 h. Senescence β-Galactosidase Staining Kit (cell signaling cat. 9860) was used to assess cellular senescence, according to the manufacturer’s instructions. Briefly, cells were rinsed in PBS and fixed with the kit provided solution for 10 min at RT. 1 ml of the β-Galactosidase staining solution was added in each well and the plate was covered with paraffin to avoid evaporation. The plate was incubated 16 h at 37 °C in a dry incubator and cells were stained with DAPI at 0.001 mg/ml for 10 min. A Nikon eclipse Ti microscope objective at 10X magnification was used for image acquisition. Pictures of 4 random representative areas were taken and the ratio of β-Galactosidase to total cells (DAPI-positive cells) was used as a readout for senescent cells.

### Statistical analyses

*In vivo* and *ex vivo* data are presented as the mean ± SD for each group. Statistical analysis was performed by non-paired (*in vivo*) or paired (*in vitro*) one-way ANOVA, followed by Tukey’s test corrections for multiple comparisons. Data in each group were checked for normal distribution using the D’Agostino and Pearson normality test based on *p* = 0.05. Human data statistical analysis was done using two-tailed T-test and spearman correlation.

## RESULT

### Deletion of SHP-1 in ECs restores blood flow and limb utilization in diabetic mice following ischemia

To study impact of SHP-1 specifically in endothelial cells following diabetic PAD, unilateral femoral artery ligation was performed in nondiabetic and diabetic ec-SHP-1^+/+^ and ec-SHP-1^-/-^ mice, which were followed for 4 weeks. Blood glucose levels and body weight of each animal were evaluated throughout the study. Notably, the absence of SHP-1 only in EC did not influence diabetes-induced elevated blood glucose and lower body weight (Supplemental Table S4). Following euthanasia, EC from lungs and SMC from aorta were harvested and digested for DNA extraction. We confirmed that SHP-1 was specifically deleted in EC and not in SMC (Supplemental Fig. S1A to S1D). To investigate blood flow reperfusion, we performed laser Doppler imaging over 28 days in the ischemic leg of all groups of mice (Fig. 1A). As previously observed, diabetic ec-SHP-1^+/+^ mice displayed 46% blood flow reperfusion following 28 days of femoral artery ligation whereas nondiabetic ec-SHP-1^+/+^ mice exhibited 81% blood flow reperfusion (*p*=0.0008; Fig.1B), representing a reduction of 43%. The specific deletion of SHP-1 in EC significantly improved blood flow of the ischemic limb at 67% (*p*=0.0293) in diabetic mice compared to diabetic mice with an ec-SHP-1^+/+^ genotype. This positive effect of endothelial SHP-1 ablation on blood flow reperfusion was associated with improved motor function of the mice. The ability to perform exercise is often reduced by diabetes due to leg pain. Thus, we assessed exercise willingness of the mice in individual cages using voluntary wheel running. Diabetic ec-SHP-1^+/+^ mice exhibited a significant 82% reduction in total distance traveled (17490 m VS 3217 m; *p=*0.0034; Fig. 1C) and number of laps (*p=*0.0014, Fig. 1D) as compared to nondiabetic counterparts. Notably, specific deletion of SHP-1 in EC showed similar motor function (15435 m VS 17490 m) as nondiabetic control mice (Fig. 1C and 1D.).

**Figure 1:**
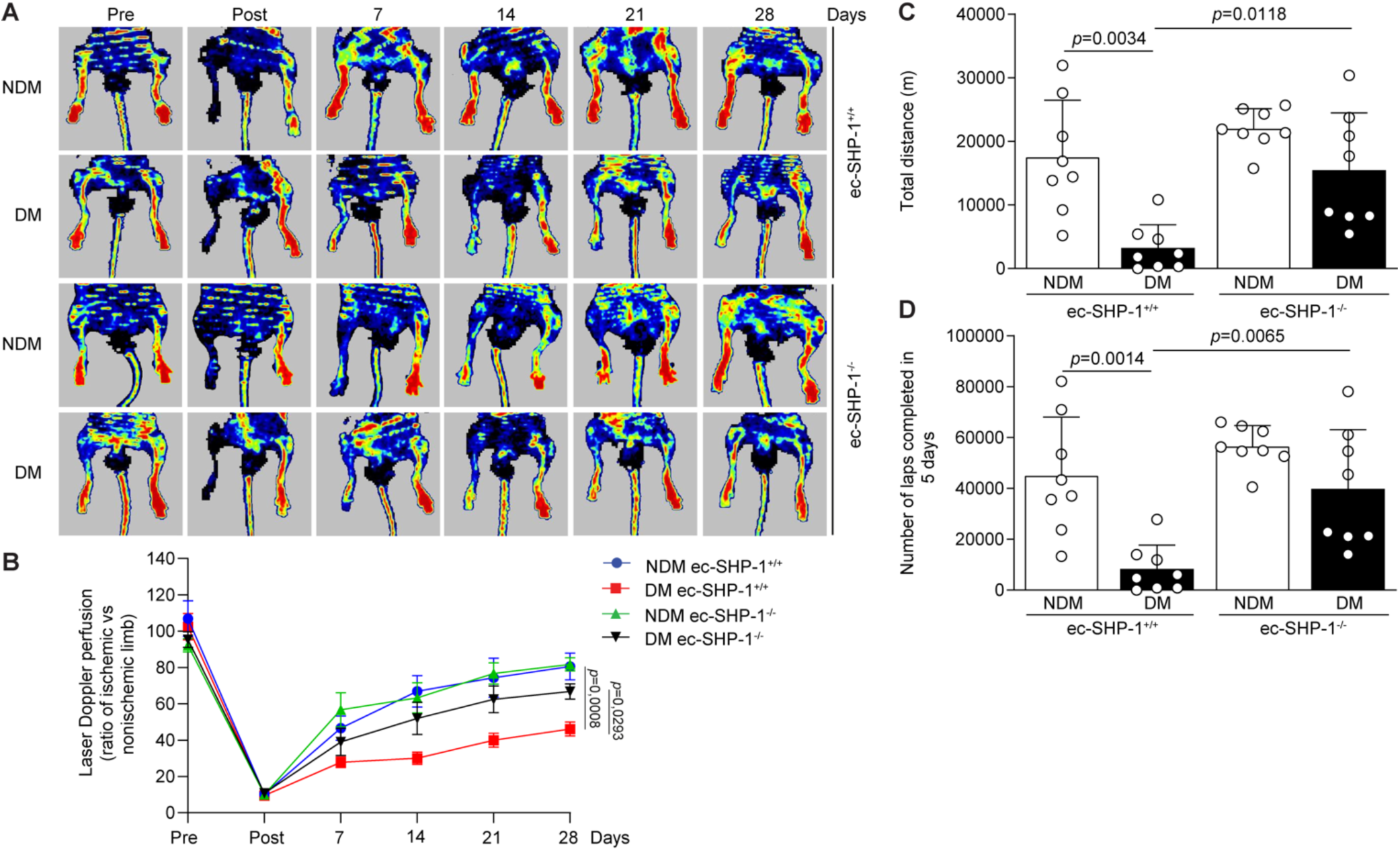
Deletion of SHP-1 in EC improves blood flow reperfusion and restores limb motor function following limb ischemia in diabetes. (A) Representative images of the laser Doopler imaging, before, immediately after, and 7,14, 21 and 28 days after femoral artery ligation. (B) Quantification of blood flow reperfusion. (C) Cumulative running distance (m) and (D) total number of completed laps over 5 days in the voluntary exercise wheel made by nondiabetic (whites bars) and diabetic mice (black bars) without (ec-SHP-1^+/+^) or with endothelial-specific SHP-1 deletion (ec-SHP-1^-/-^). Results are shown as mean ± SD of 8-9 mice per group. One-way ANOVA with Tukey’s *post hoc* test.

### Endothelial-specific deletion of SHP-1 prevents diabetes-induced myofiber atrophy and improves vessel density

To evaluate the structure of muscle fibers in our animal models we measured the mean myofiber diameter (Fig. 2A). As previously observed, diabetic wild-type mice displayed lower mean myofiber diameter compared to nondiabetic wild-type mice (*p<*0.0001; Fig. 2B), this reduction was prevented by endothelial-specific deletion of SHP-1 (*p=*0.0043; Fig. 2B). Diabetes is associated with impaired blood flow reperfusion and angiogenesis. Indeed, our data confirmed that three months of uncontrolled diabetes blunted small vessel density by reducing the number of capillaries by 42% (*p=*0.0089; Fig. 2C, 2D) and the number of arterioles by 56% (*p=*0.0089 Fig. 2C, 2E) compared to nondiabetic ec-SHP-1^+/+^ mice. Ablation of SHP-1 specifically in EC was able to restore small vessel density in diabetic mice to the level of nondiabetic controls (*p=0.0023* Fig. 2C, 2D, and *p=<*0.0001; 2E).

**Figure 2:**
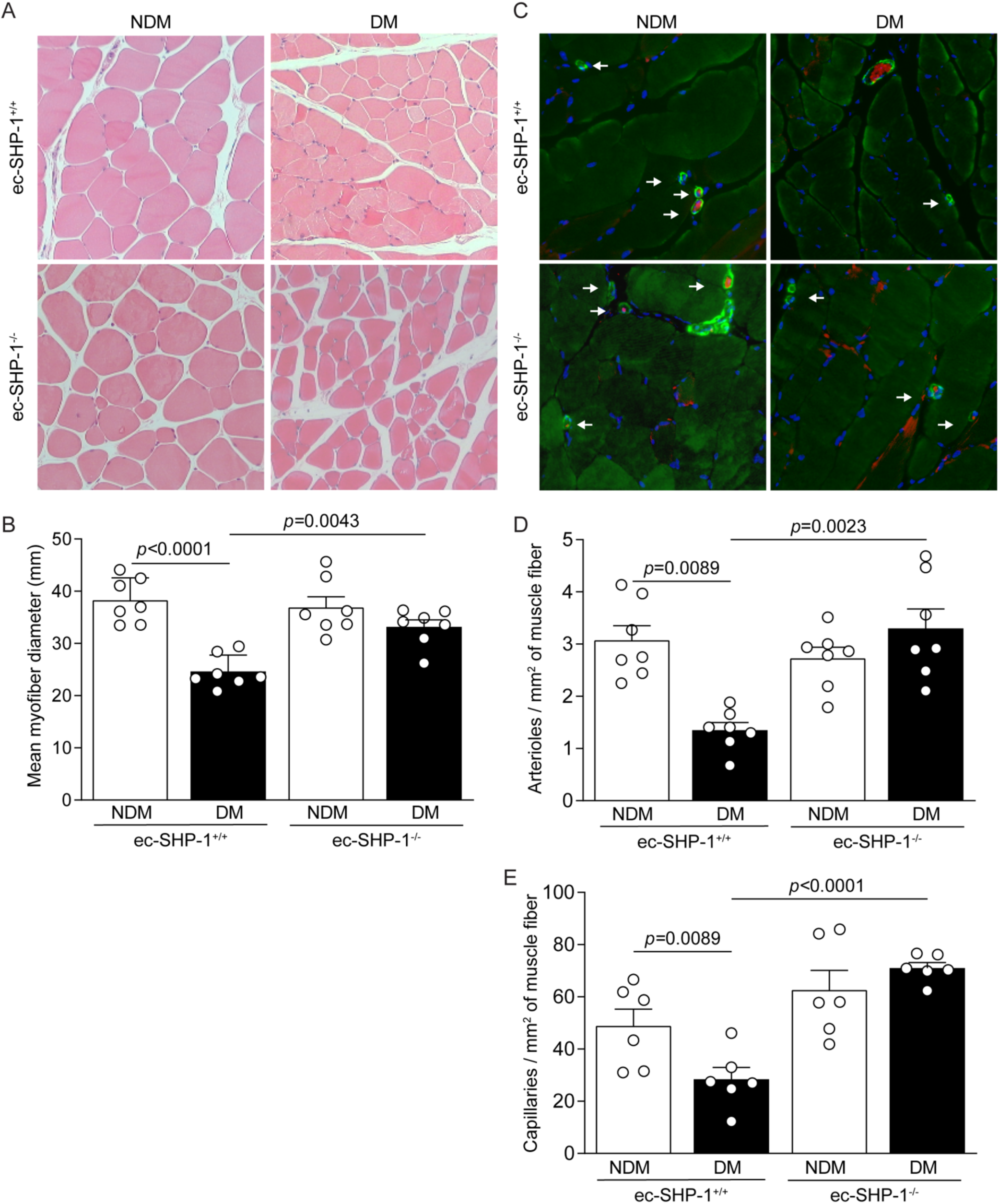
Endothelial specific ablation of SHP-1 maintains muscle fiber integrity and vascular density. (A) Morphological analysis showing ischemic muscle stained with hematoxylin and eosin (H&E). (B) Structural analysis and quantification of the mean myofiber diameter of the ischemic muscle. (C). Immunofluorescence of endothelial cells in red, smooth muscles cells in green and nuclear cell (DAPI) in blue on ischemic abductor muscle cross section. (D) Quantification of the number of arterioles and (E) capillaries of the entire muscle cross section reported by the muscle fiber area. Mice groups are presented as nondiabetic (white bars) and diabetic mice (black bars) without (ec-SHP-1^+/+^) or with endothelial-specific SHP-1 deletion (ec-SHP-1^-/-^). Results are shown as mean ± SD of 6 to 7 mice per group. One-way ANOVA with Tukey’s *post hoc* test.

### Endothelial-specific SHP-1 deletion restores angiogenesis and prevents senescence in diabetic mice

We have previously reported that diabetes-induced SHP-1 in ischemic muscle is associated with decreased VEGF receptor activation^17^. However, direct in vivo evidence that SHP-1 specifically in endothelial cells is responsible for VEGF inhibition is lacking. Our data confirmed that reduced VEGFR2 and Akt phosphorylation in diabetes can be prevented by SHP-1 deletion in endothelial cells (*p*=0.0014 Fig. 3A). In addition, phosphorylation (*p*=0.0149 Fig. 3B) and expression (*p*=0.0011 Fig. 3C) of eNOS, a well-known target of VEGF and an important protein for EC function, was decreased in ischemic muscle of diabetic mice. Endothelial-specific deletion of SHP-1 restored both expression and phosphorylation of eNOS in diabetic mice. Moreover, mRNA expression analysis in EC of diabetic mice showed decreased expression of Flk-1 and VEGF (*p*=0.0156 Fig. 3D and *p*=0.0216 Fig. 3E), which was not observed if SHP-1 was deleted in EC. Because cellular senescence is involved in vascular and metabolic disorders such as cardiovascular disease, obesity and diabetes^21^, we measured senescence markers in the ischemic muscle. Our data revealed that expression of the tumor suppressor p53 and the cyclin-dependent kinase inhibitor p21 was upregulated whereas NRF-2, an antioxidant factor associated with delayed cell senescence, was downregulated in ischemic muscle of diabetic ec-SHP-1^+/+^ mice compared to nondiabetic littermate controls (Fig. 3F, 3G and 3H). Interestingly, deletion of SHP-1 in endothelial cells reversed diabetes-induced p53 and p21 expression and restored NRF-2 expression in ischemic muscle (Fig. 3F, 3G and 3H).

**Figure 3:**
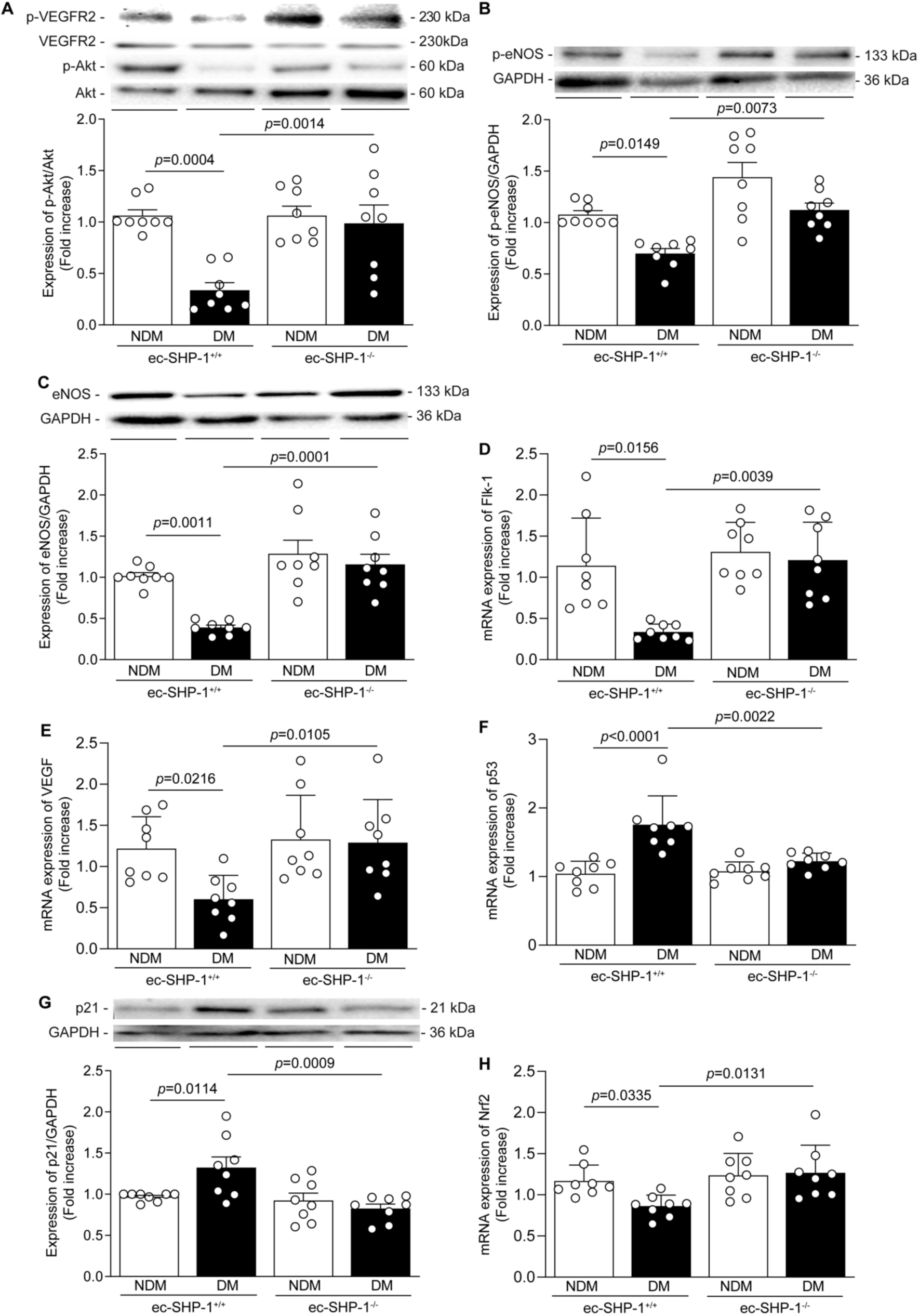
Endothelial cell dysfunction and senescence caused by diabetes are prevented by the inhibition of SHP-1. Immunoblot and quantification of (A) VEGFR2, p-Akt, (B) p-eNOS, (C) eNOS, (G) p21 protein expression and mRNA expression of (D) Flk-1, (E) VEGF-a, (F) p53 and (H) Nrf2 in nondiabetic (whites bars) and diabetic mice (black bars) without (ec-SHP-1^+/+^) or with endothelial-specific SHP-1 deletion (ec-SHP-1^-/-^). Results are shown as mean ± SD of 8 mice per group. One-way ANOVA with Tukey’s *post hoc* test.

### High glucose-induced SHP-1 blocks VEGF induced angiogenesis and enhances EC senescence

Our *in vivo* data suggests that SHP-1 has negative effects on endothelial function in response to hyperglycemia and ischemia. Thus, we performed additional experiments in cultured EC to investigate its mechanisms. As previously reported by us and others, exposure to high glucose (HG) levels inhibits the pro-angiogenic action of VEGF in EC^19,29^. In line with these observations, we observed a significant reduction of EC proliferation (Fig. 4A), migration (Fig. 4B and 4C) and tubule formation (Fig. 4D, 4E) under HG conditions *in vitro*. Importantly, overexpression of adenoviral of dominant-negative form of SHP-1 was able to completely re-establish the angiogenic effects of VEGF on proliferation, migration and tubule formation under HG conditions (Fig. 4A to 4E). In high glucose conditions, SHP-1 protein is bound to VEGFR2 receptors (Supplemental Fig. 2A), which can prevent VEGF-induced the phosphorylation of its receptor and downstream target Akt (Supplemental Fig. S2B and S2C). Restoring VEGF actions with the overexpression of SHP-1 dominant negative adenovirus was led to elevated phosphorylation of VEGRF2 and Akt (Supplemental Fig. S2B and S2C). Beside blocking the effects of VEGF, HG exposure enhanced EC senescence, measured by the presence of SA β-galactosidase positive cells (*p*=0.0219; Fig. 5A and 5B). In addition, EC exposed to HG showed increased expression of p21 (*p*=0.0342 Fig. 5C) and decreased expression of Nrf2 (*p*=0.0427 Fig. 5D). Interestingly, overexpression of a dominant negative form of SHP-1 prevented HG induced endothelial cell senescence and cell cycle arrest (*p*=0.0009 Fig. 5C and *p*=0.0225 Fig. 5D).

**Figure 4:**
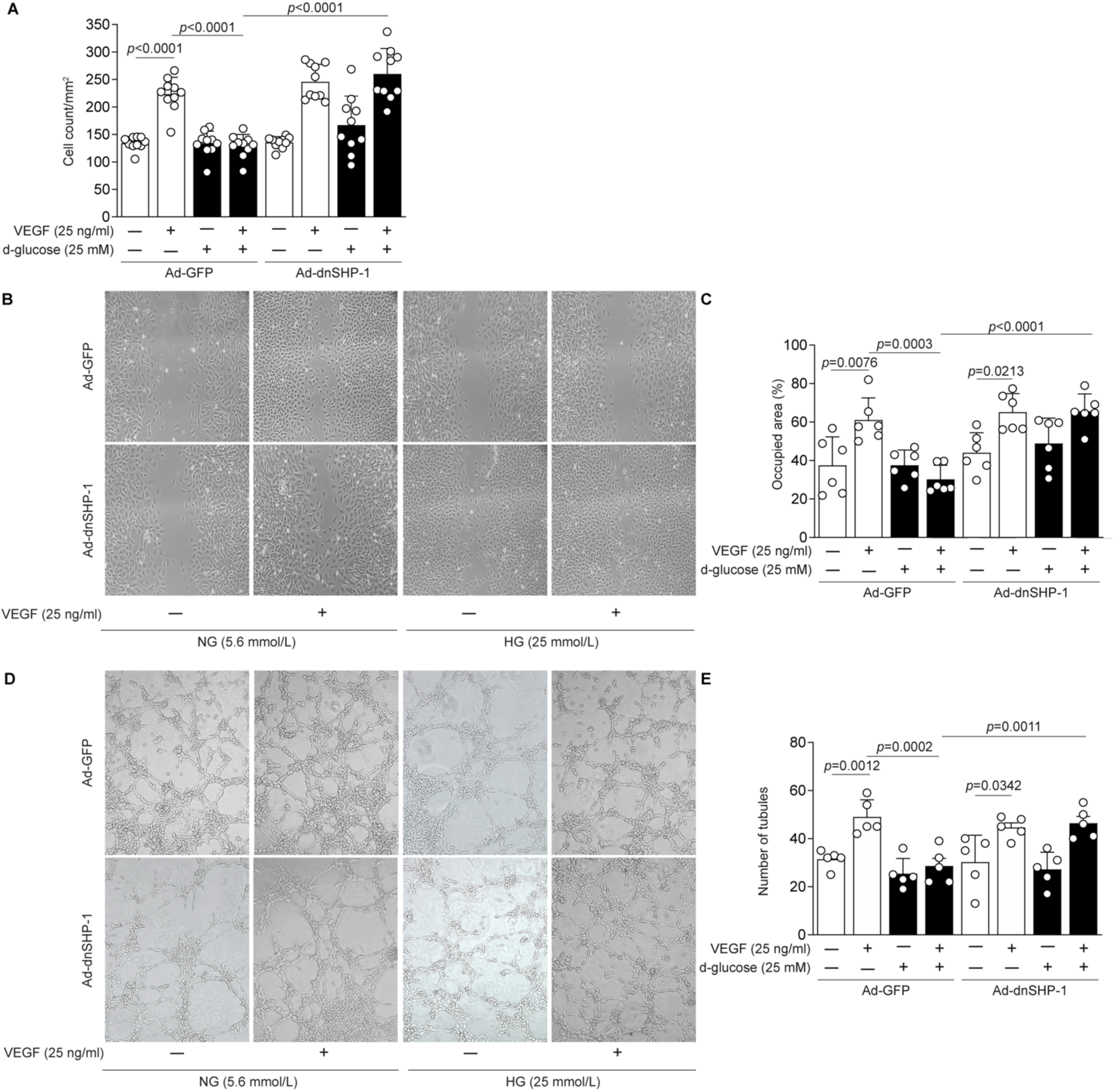
Inhibition of SHP-1 prevents HG-induced reduction of VEGF actions in EC. Cultured EC were infected with adenoviral GFP or SHP-1 dominant-negative form and then incubated with normal glucose (NG; 5.6 mmol/L; white bars) or high glucose (HG; 25 mmol/L; black bars) and then stimulated with VEGF-A for 16h (A-C), 4h (D-E) under hypoxia (1% O2) for the last 16h of treatment in all experiments. (A) Quantification of cells count/mm^2^. (B) Representative images of the EC migration assay. (C) The percentage of the surface area occupied by the EC was quantified. (D) Representative images of the lumen formation abilities of EC. (E) Tubule formation was quantified by measuring the total number of closed circles in the entire well, normalized on the NG condition. Results are shown as mean ± SD of 5 to 10 independent cell replicates. One-way ANOVA with Tukey’s *post hoc* test.

**Figure 5:**
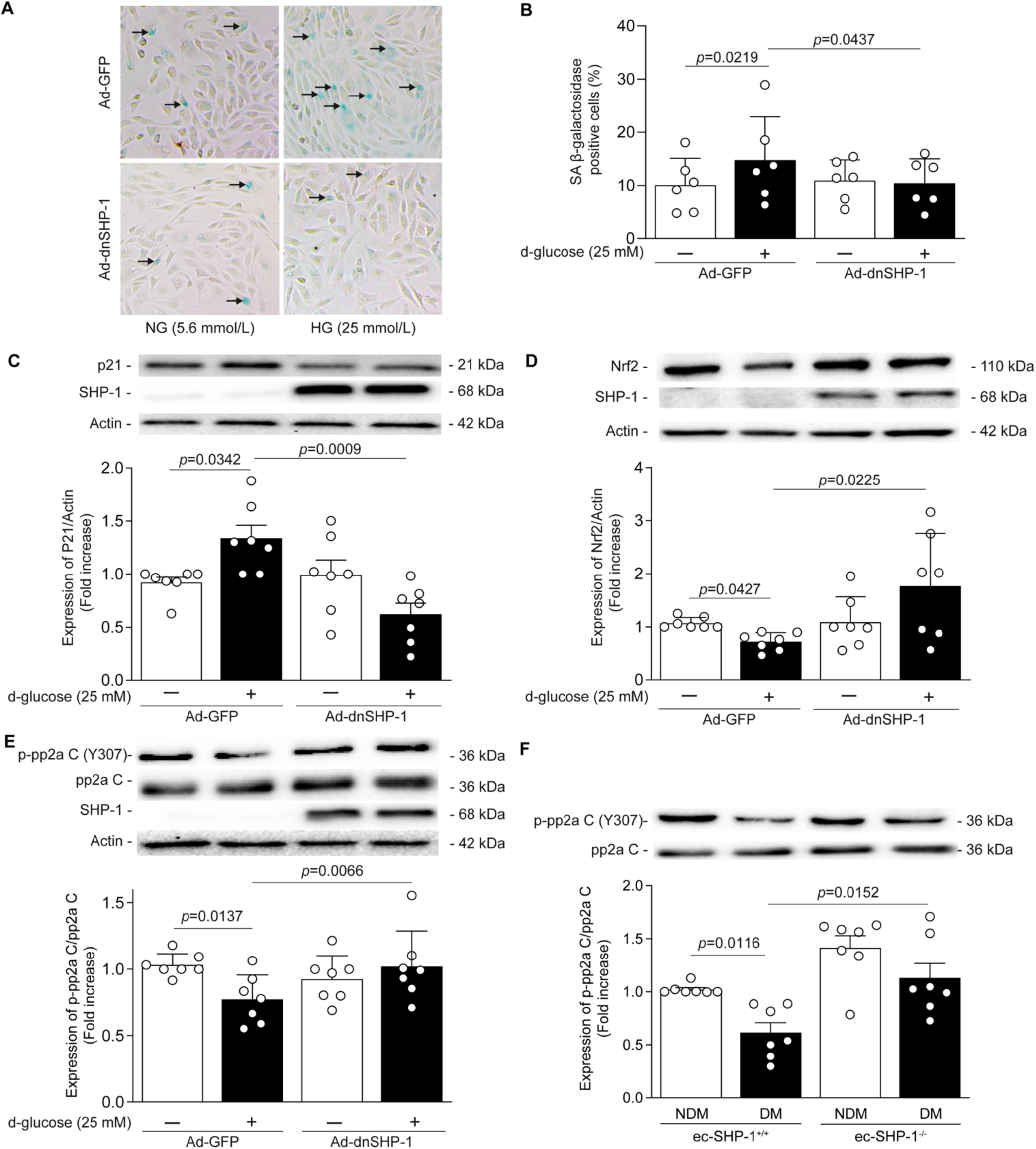
SHP-1 regulates endothelial senescence under HG level exposure. (A) Representative images of senescence-associated β-galactosidase assay and (B) quantification of the number of positive cells (blue staining). Immunoblot representative image and quantification of (C) p21, (D) Nrf2 and (E-F) phospho-pp2aC (Y307). EC were infected with adenoviral GFP or SHP-1 dominant-negative form and exposed to normal glucose (NG; 5.6 mmol/L; white bars) or high glucose (HG; 25 mmol/L; black bars) as well as in (F) nondiabetic (white bars) and diabetic (black bars) mice without (ec-SHP-1^+/+^) or with endothelial-specific SHP-1 deletion (ec-SHP-1^-/-^). Results are shown as mean ± SD of 7 independent cell replicates or mice per group. One-way ANOVA with Tukey’s *post hoc* test.

### SHP-1 regulates p53/p21 through p-PP2a dephosphorylation

Previous studies have shown that a specific subunit of the serine/threonine phosphatase PP2A becomes activated to support the activity of p53 upon metabolic stress^30,31^. We observed that EC exposed to HG exhibited a significant reduction of PP2a C subunit activity measured by decreased tyrosine Y307 phosphorylation (*p*=0.0137; Fig. 5E). However, loss of tyrosine Y307 phosphorylation in EC exposed to HG was prevented by the overexpression of dominant negative SHP-1 (*p*=0.0066; Fig. 5E). This finding was confirmed by a decrease tyrosine 307 phosphorylation in ischemic muscle of diabetic mice, which was counteracted by EC-specific deletion of SHP-1 (*p*=0.0152; Fig. 5F).

### SHP-1 and EC senescence in human patients with PAD

To support our preclinical findings in the context of human biology, tibial artery and gastrocnemius muscle from patients without or with diabetes that underwent lower limb amputation were analysed. We observed an increase in the expression of SHP-1 in gastrocnemius muscle of patients with diabetes and PAD compared to patients with PAD but without diabetes (*p*=0.022; Fig. 6A). Patients with diabetes also exhibited higher levels of p53 (*p*=0.0165, Fig. 6B) p21 (*p*=0.0383, Fig. 3C) mRNA. Interestingly, SHP-1 expression was also elevated in tibial artery of patients with diabetes and PAD compared to those without diabetes (*p*=0.0338; Fig. 6D). Although p53 expression was not significantly elevated in tibial artery with diabetes (Fig. 6E), there is strong correlation between elevated levels of SHP-1 and p53 expression (ρ*=*0.633*; p*=0.0038, Fig 6F). In addition, we observed a significant 44% reduction in Nrf2 mRNA levels in tibial artery with diabetes (*p*=0.0138, Fig. 6G) and SHP-1 expression correlated negatively with Nrf2 (ρ=-0.350*; p*=0.0173; Fig. 6H).

**Figure 6:**
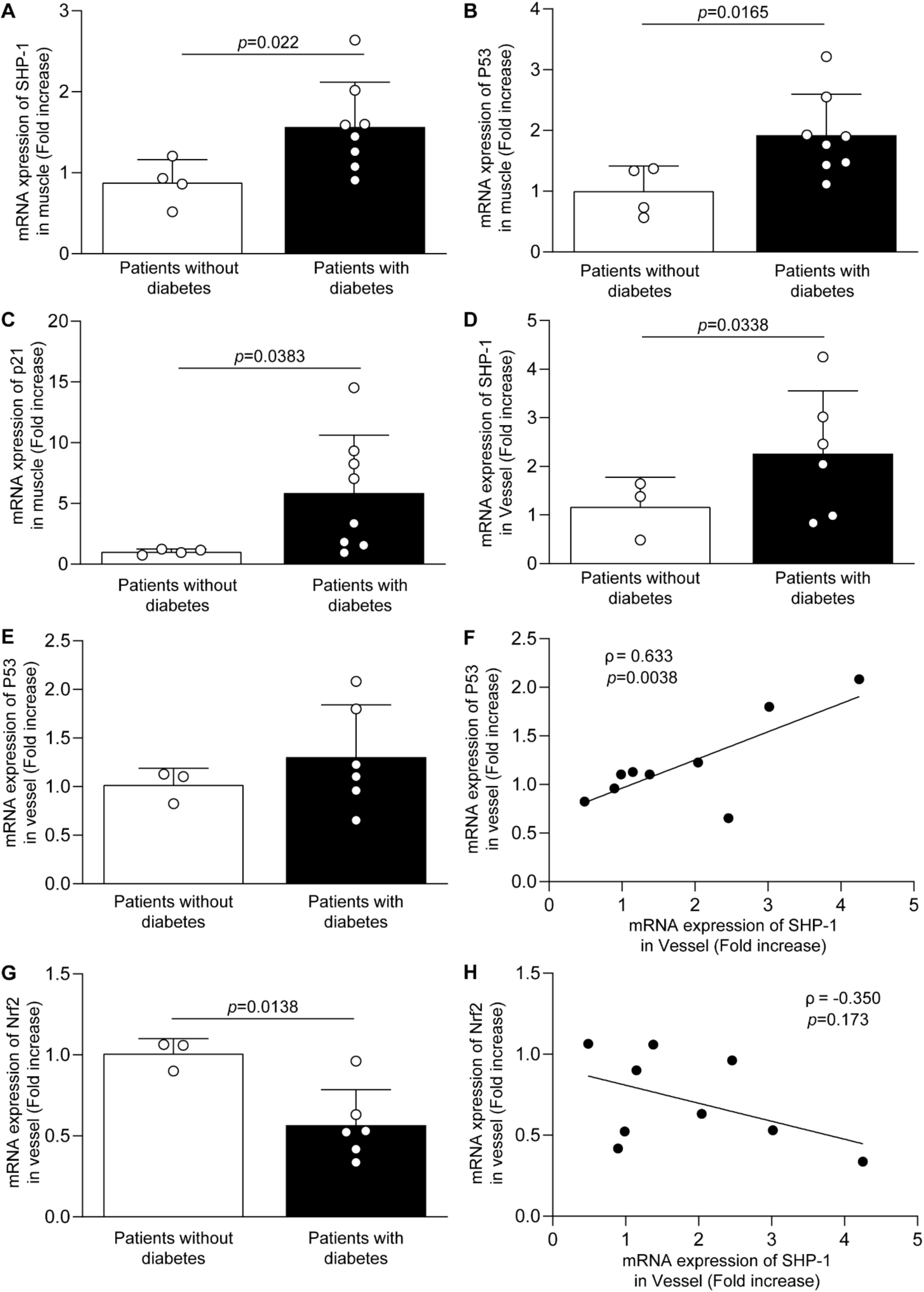
Elevated expression of SHP-1 in ischemic muscle and artery of patients with diabetes is associated with cellular senescence. (A,D) SHP-1, (B, E) p53, (C) p21 and (G) Nrf2 mRNA expression in (A-C) gastrocnemius muscle and (D-H) tibial vessels of patients without (white bars) and with diabetes (black bars) that suffered from an amputation. Correlation between SHP-1 expression and (F) p53 and (H) Nrf2. Results are shown as mean ± SD (A-E, G) One-tailed t test or Spearman correlation (F and H) of 3 to 8 patients per group.

## DISCUSSION

With a growing diabetes epidemic and an aging patient population in modern countries, the prevalence of PAD will continue to rise in the near future^32^. Although new classes of drugs (SGLT2 inhibitors and GLP-1 agonists) have improved glycemic control and cardiometabolic health, there is still no effective treatment that reduce the incidence of PAD in patients with diabetes^1^. Unfortunately, even the benefits of supervised and home-based walking exercises on mobility in patients with PAD are dampened by poor adherence and lack of durability^33^. Therefore, a better understanding of mechanisms of diabetes-induced abnormal collateral vessel formation and muscle damage in PAD may help the identification of new therapeutic options for this population.

It has been more than two decades since Rivard and colleagues demonstrated that blood reperfusion following ischemia was impaired in diabetic mice compared to nondiabetic mice, mainly due to a reduction in VEGF signaling^34^. We previously demonstrated that diabetes increases the expression and activity of SHP-1 in ischemic muscles and EC through protein kinase C delta^17^. However, direct evidence of an involvement of SHP-1 in endothelial cell function has never been explored. Results of the present study show a significant blood flow reperfusion recovery 4 weeks after surgery in the ischemic muscle of diabetes mice that do not express SHP-1 specifically in EC compared to diabetic wild-type mice. The improvement in blood flow reperfusion is accompanied by an improvement in motor function compared to diabetic mice. Interestingly, EC-specific deletion of SHP-1 did not restore blood flow reperfusion to the same level as the nondiabetic ec-SHP-1+/+ mice, suggesting that deletion of SHP-1 in endothelial cells alone is not sufficient to fully prevent the impact of diabetes on angiogenic processes. Indeed, our group has reported that SHP-1 is also implicated in smooth muscle cell function^16^. Moreover, deletion of SHP-1 in EC did not restore the expression of PDGF, an important growth factor for vessel maturation, and its receptor in ischemic muscles.

Previous animal and human studies in patients without and with diabetes in revascularization following ischemia have reported decreased expression of different pro-angiogenic factors, including VEGF-A, VEGFR-2, PDGF-B, PDGFR-β and HIF-1α in the diseased population^35^. VEGF is one of the main regulators of angiogenesis, activating different EC function such as cell migration, proliferation, and survival to enable the formation of new vessels and restore oxygenation of the limb following ischemia. The VEGF signaling pathway requires the activation of its receptor, VEGFR-2, to induce several signaling cascades that activate various downstream processes^36^. We have previously shown that the increased expression and phosphatase activity of the SHP-1 protein correlates with a reduction of VEGFR-2 phosphorylation in diabetic ischemic muscle^17^. The present study confirmed that elevated SHP-1 protein is binding to VEGFR-2 under HG conditions. This interaction is associated with a significant decrease in EC proliferation, migration, tubule formation and eNOS expression and activity under HG levels. It has also been shown that inhibition of SHP-1 by siRNA in endothelial cells increased phosphorylation of tyrosine residues of VEGFR2, increasing cell proliferation, but not cell migration^14,37^. Our study provides evidence that overexpression of an inactive form of SHP-1 in endothelial cells exposed to HG prevented the EC dysfunction caused by diabetes, indicating that SHP-1 directly inhibits VEGFR-2 activation. In addition, we also provide *in vivo* confirmation that deletion of SHP-1 in EC prevented diabetes-induced aberrant collateral vessel formation, motricity impairment, and endothelial cell apoptosis by restoring muscle integrity and vascular density.

Senescence is a process of cell cycle arrest associated with cellular aging. Vascular senescence is associated with several cardiovascular pathologies, including PAD^24^. Exposure of human aortic EC to serum from patients with PAD caused an increase in the expression of p21 and p53, as well as activity of senescence-associated β-galactosidase^38^. Moreover, streptozotocin-induced diabetes caused an increase in p53 hyperacetylation in the aorta compared to control mice^39^, suggesting that diabetes may activate the senescence process of vascular cells. In a mouse model of cardiac fibrosis, accumulation of p53 in endothelial cells lead to vessel rarefaction and inhibition of endothelial sprouting vessel formation, suggesting the importance of senescence in endothelial function under ischemia conditions^40^. Our study provides novel evidence that p53 and p21 are elevated in ischemic in muscle and vessels of patients with diabetes and PAD and EC-specific deletion of SHP-1 blocked p53 and p21 expression in diabetic ischemic muscle. A previous study by Sun et al. investigated a role of SHP-1 in nasopharyngeal carcinoma cell senescence. In contrast to our observations, decreased SHP-1 expression was associated with senescence activation, due to increased expression of p16 and pRb. However, Sun et al. did not detect any changes in p53 and p21 expression^41^. Moreover, it is likely that cancer cells react differently than vascular cells. Yokoi and colleagues reported that endothelial cell exposure to HG enhanced the number of senescent cells compared to cells exposed to normal glucose levels^42^. Our data confirmed this observation and provided novel insights that inhibition of SHP-1 is able to prevent EC senescence, p53 and p21 expression induced by HG. Nrf2, a redox-sensitive transcription factor, is expressed in all cell types and regulates antioxidant defense and cell processes such as mitochondrial bioenergetics, autophagy, unfolded protein response and metabolism through the activation of cytoprotective genes^43^. Previous work has shown that the activation of Nrf2 pathway promotes antioxidant signals and inhibits cellular senescence, thereby exerting protective effects against age-associated disease^44^. Furthermore, Nrf2 activation in diabetic ischemic muscle has been linked to improved revascularization in a mouse model of type 2 diabetes^45^. Our data demonstrate that decreased expression of Nrf2 is observed in the ischemic muscle of diabetic ec-SHP-1^+/+^ mice compared to NDM ec-SHP-1^+/+^ mice, whereas endothelial-specific deletion of SHP-1 increased Nrf2 expression in diabetic ischemic muscle. In addition, a correlation between Nrf2 and p53/p21 in response to oxidative stress has been reported^46,47^.

Our results also support the idea that diabetes-induced SHP-1 activation leads to enhanced p53 and p21 expression and Nrf2 degradation in mice and patients with diabetic ischemic muscle. Our *in vitro* results are consistent with our *in vivo* results, showing an increase in p21 expression and the number of senescent EC, quantified using senescence-associated β-galactosidase labeling, in addition to a decrease in Nrf2 expression in EC exposed to HG, which are effects that were prevented by overexpression of a dominant negative form of SHP-1. These results demonstrate an involvement of SHP-1 in the activation of EC senescence due to activation of the p53/p21 pathway and the inhibition of Nrf2.

Increased serine/threonine phosphatase PP2A activity has previously been associated with increased p53 activity^30,31^. Phosphorylation of threonine 55 of p53 inhibits its activity. Vice-versa dephosphorylation of threonine 55 from p53 by PP2A induces its activation and senescence processes^30^. The activity of PP2A is regulated by the phosphorylation of its tyrosine 307 (Y307). Phosphorylation of this residue inhibits the binding of the PP2A “B” subunit, preventing its catalytic activity^48^. Therefore, we assessed levels of PP2A phosphorylation in diabetic ischemic muscle as well as in EC exposed to a HG under hypoxia conditions. We observed a significant decrease in tyrosine 307 phosphorylation of PP2A in ischemic muscle of DM ec-SHP-1^+/+^ mice compared to NDM ec-SHP-1^+/+^ mice as well as in EC under HG conditions. These results are consistent with a study by Du and colleagues that also observed an increase in PP2A activity in EC exposed to high glucose concentration, causing elevated oxidative stress and cell death^49^. Endothelial-specific deletion of SHP-1 in diabetic ischemic muscle and the overexpression of the negative dominant adenoviral vector of SHP-1 prevented the activation of PP2A, suggesting a relationship between SHP-1 and activation of PP2A and a potential explanation of SHP-1 involvement in the regulation of EC senescence in diabetic ischemic muscle.

In conclusion, our present study provided further insights on the mechanisms caused by diabetes that are associated with EC dysfunction and senescence. In addition, our study provides novel mechanistic pathways regulated by SHP-1 in the context of diabetes and poor collateral vessel formation that are associated with endothelial cell damage and dysfunction.

## Acknowledgements

The authors gratefully acknowledge Marilène Paquette (Histology Core, U. of Sherbrooke) for her assistance. Dr. Geraldes is the guarantor of this work, has full access to all the data, and takes full responsibility for the integrity of the data and the accuracy of data analysis. This work was supported by grants from the Canadian Institute of Health Research [PJT153165 to P.G.] This work was performed at the CHUS research center, funded by the “Fonds de Recherche du Québec – Santé” (FRQS). Dr. Geraldes is the holder of the Canada Research Chair in Vascular Complications of Diabetes. C.F.B. was supported by the Canadian Institutes of Health Research (CIHR, PJT-162442), the Natural Sciences and Engineering Research Council of Canada (NSERC, RGPIN-2017-05490), the Fonds de Recherche du Québec - Santé (FRQS, Dossiers 296357, 34813, and 36789), the Fonds de Recherche du Québec - Nature et Technologies (FRQNT, Dossier 331297), and the Canadian Stem Cell Network.

## Author Contributions

Author contributions: A.N., M.O., R.B, C.M., and S.R. performed experiments, sample and data collection, and experimental data. F.B. provided the Cdh5-CreERT2 mice and edited the manuscript. F.L. and M-A.D recruited patients, process and analyzed human tissues samples. A.N., F.L. and P.G. analyzed the data and wrote the manuscript.

## Supplemental Material and methods

### Mice genotyping

Kidney glomeruli were digested with proteinase K O/N at 55 °C in final concentration of 1X Modified Gitschier Buffer (Tris HCL pH 8.8 67 mM, Amonium Sulfate 16 mM, MgCl2 6.5 mM) with 0.5% triton and 1% 2-beta-mercaptoethano. Presence of *loxP* and transgene were validated with specific primers listed in Supplementary Table S3. After doxycycline treatment, recombination of the *loxP* was validated using the same forward (F1) SHP-1 primer which target the 3’ lox P, and a second reverse primer (R2) which target is located after the second *loxP* sequence (see genotyping strategy in Supplementary Figure S1).

**Supplemental Table S1:** Human characteristics.

**1.1 Patients with muscle biopsy**
|  | NDM | DM | *p value |
| --- | --- | --- | --- |
| nb of patient | (n=4) | (n=8) |  |
| Gender F/M | 1/3 | 2/6 | 0,9999 |
| Age (years) | 50 ± 13 | 71 ± 8 | 0,0171 |
| Active smoking | 0 | 1 (11%) | 0,9999 |
| BMI (Kg/m<sup>2</sup>) | 24 ± 2 | 29 ± 6 | 0,03677 |
| HbA1C (%) | 5,3 ± 0,1 | 8,2 ± 0,9 | 0,002 |
| Years of diabetes | 0 | 20 ± 11 | 0,0061 |
| SHP-1 mRNA levels | 0,879 ± 0,28 | 1,567 ± 0,55 | 0,022 |

**1.2 Patients with tibial artery**
|  | NDM | DM | *p value |
| --- | --- | --- | --- |
| nb of patient | (n=3) | (n=6) |  |
| Gender F/M | 1/3 | 1/5 | 0,9999 |
| Age (years) | 63 ± 17 | 68 ± 2 | 0,869 |
| Active smoking | 0 | 1 (11%) | 0,9999 |
| BMI (Kg/m<sup>2</sup>) | 22 ± 2 | 34 ± 9 | 0,2619 |
| HbA1C (%) | 5,1 ± 0,2 | 8,2 ± 0,3 | 0,0119 |
| Years of diabetes | 0 | 36 ± 11 | 0,0119 |
| SHP-1 mRNA levels | 1,00 ± 0,46 | 2,27 ± 1,27 | 0,0338 |

**Supplemental Table S2:**
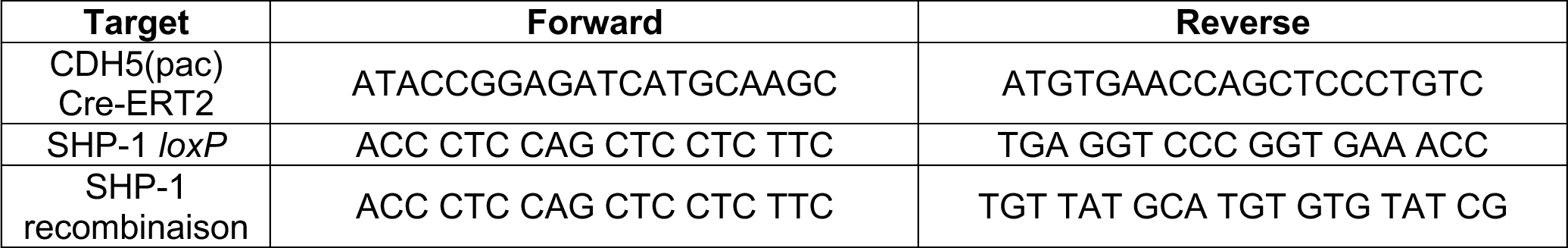
Primer sequence used for genotyping and SHP-1 specific deletion validation.

**Supplemental Table S3:**
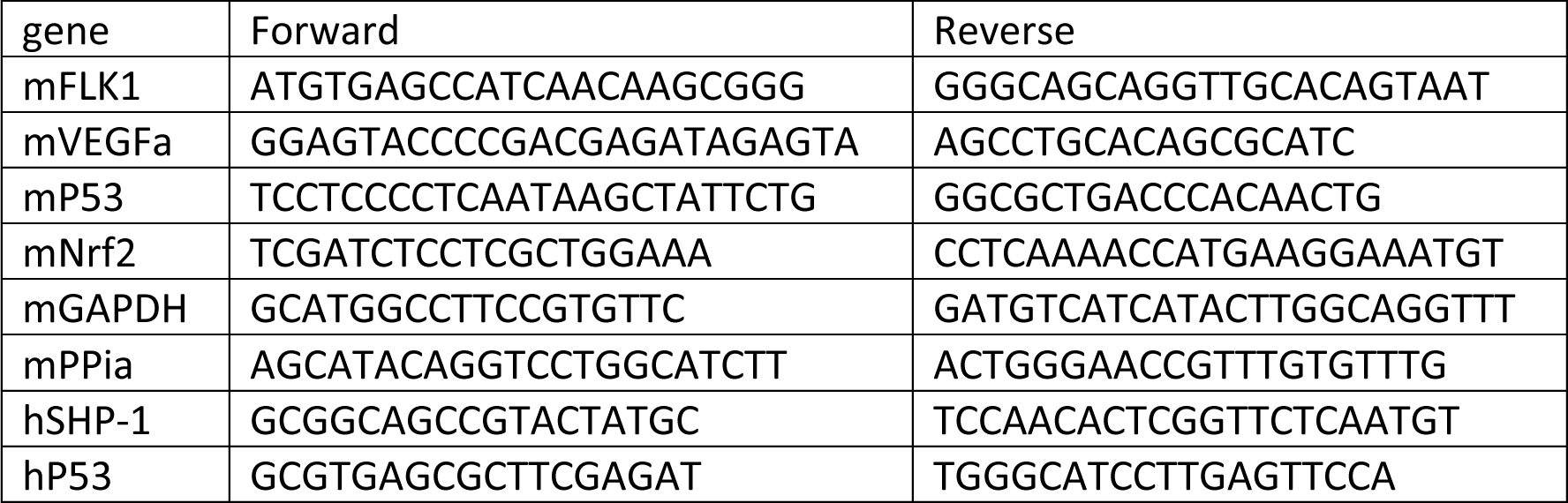
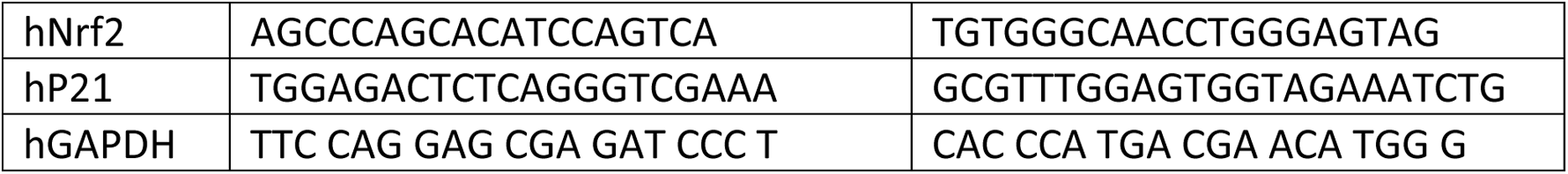
mouse (m) and human (h) primer sequence used for qPCR.

**Supplemental Table S4:**
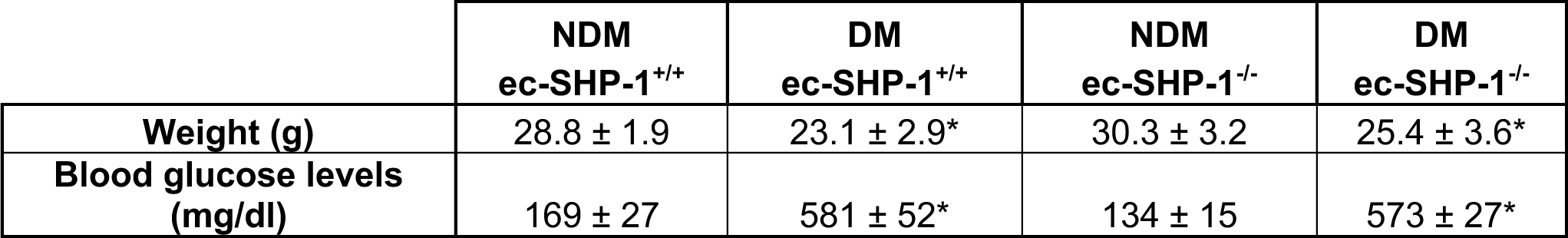
body weight and glucose levels.

**Supplemental Figure S1:**
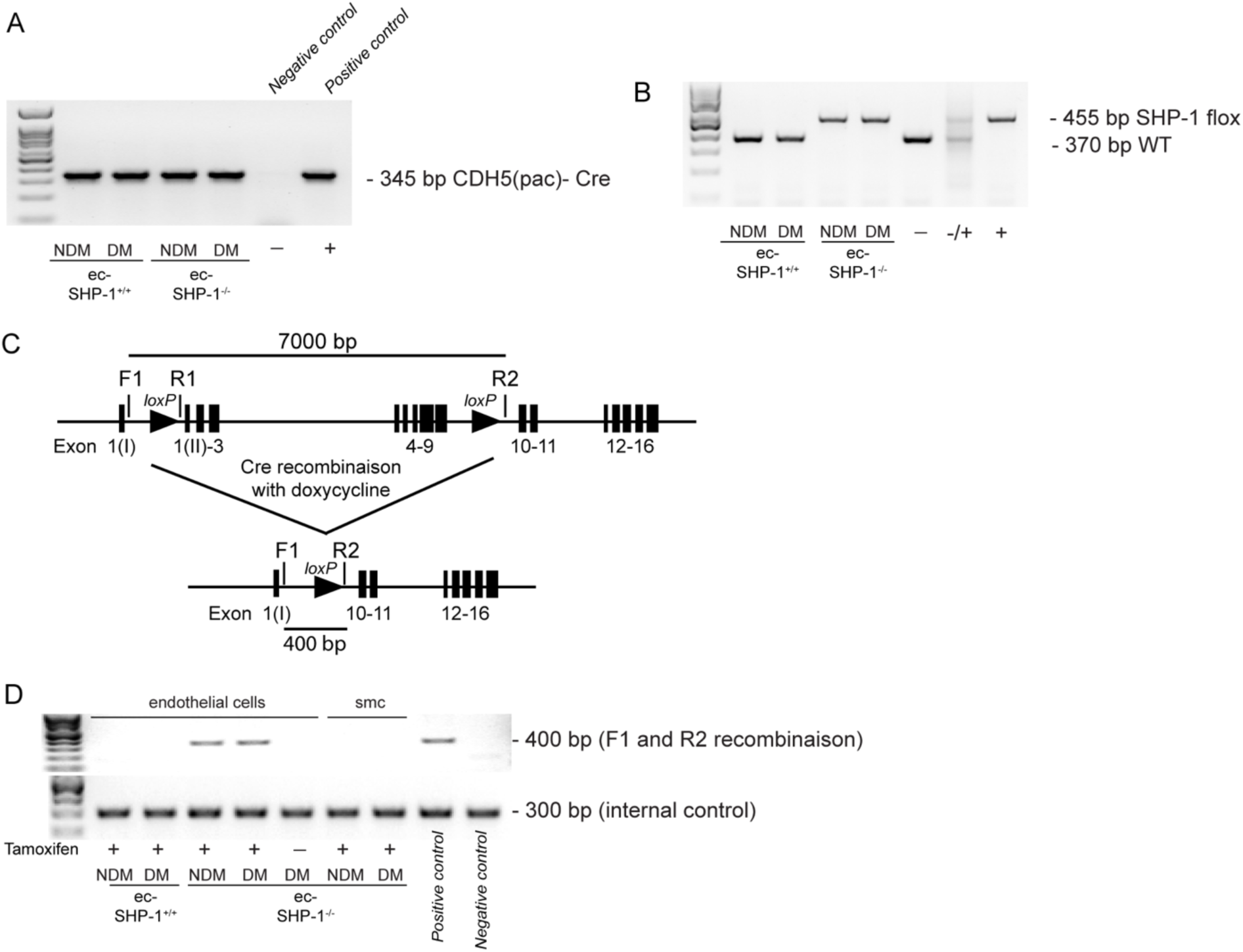
(A-B) Genotyping strategy and (C-D) confirmation of the recombination and deletion of SHP-1 specifically in endothelial cells.

**Supplemental Figure S2:**
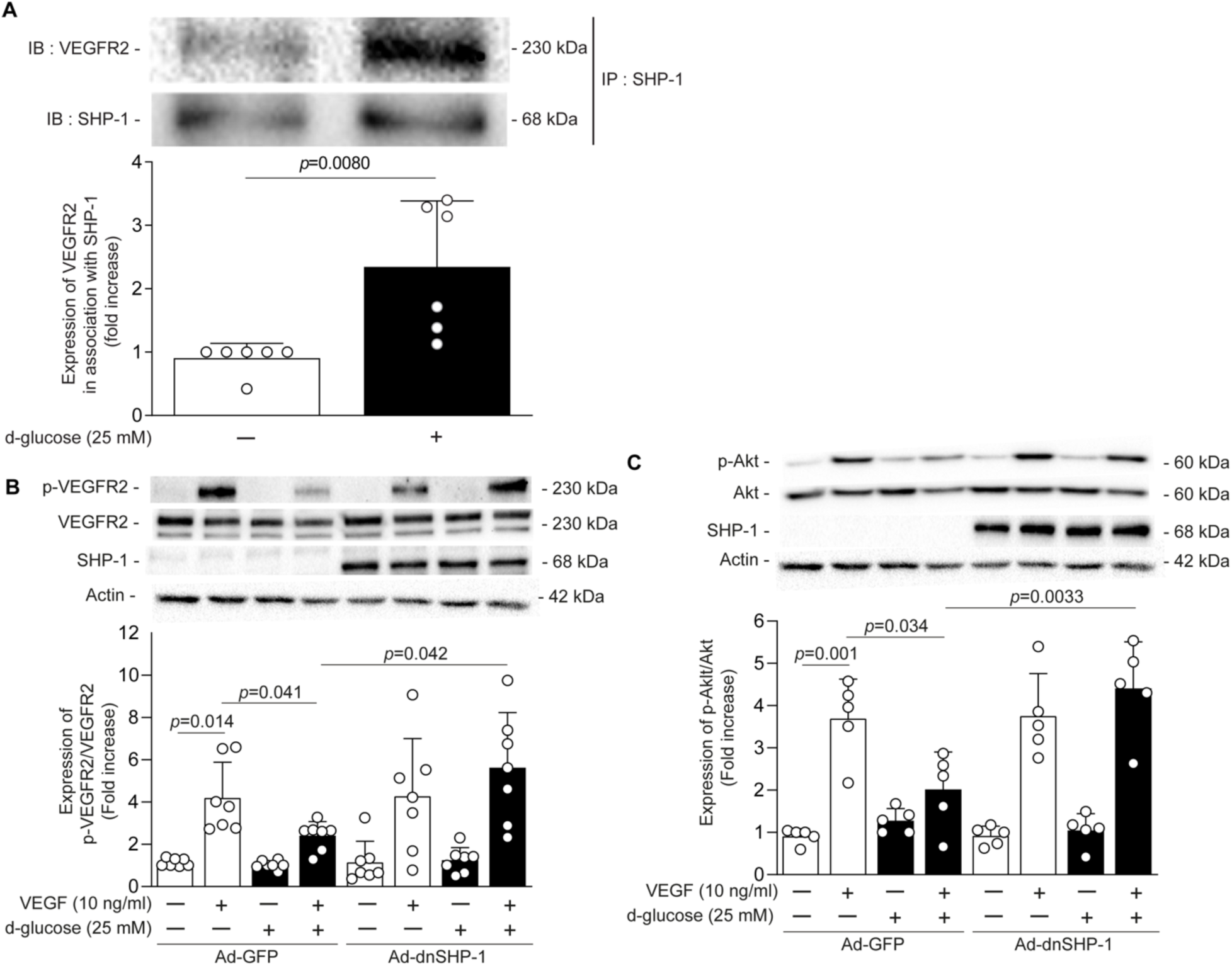
Dominant negative of SHP-1 restores VEGF signaling pathways in EC exposed to HG levels. (A) Immunoprecipitation assays of SHP-1 interaction with VEGFR2. Immunoblot representative image and quantification of (C) VEGFR2 and (C) Akt phosphorylation in EC infected with adenoviral GFP or SHP-1 dominant-negative form that were exposed to normal glucose (NG; 5.6 mmol/L; white bars) or high glucose (HG; 25 mmol/L; black bars) and then stimulated with VEGF-A for 5 min (B-C) under hypoxia (1% O2) for the last 16h of treatment in all experiments. Results are shown as mean ± SD of 5-6 independent cell replicates. One-way ANOVA with Tukey’s *post hoc* test.

## References

1. Polonsky TS, McDermott MM. Lower Extremity Peripheral Artery Disease Without Chronic Limb-Threatening Ischemia: A Review. JAMA. 2021;325:2188–2198. doi: 10.1001/jama.2021.2126

2. Hussain MA, Al-Omran M, Salata K, Sivaswamy A, Forbes TL, Sattar N, Aljabri B, Kayssi A, Verma S, de Mestral C. Population-based secular trends in lower-extremity amputation for diabetes and peripheral artery disease. CMAJ. 2019;191:E955–E961. doi: 10.1503/cmaj.190134

3. Sutton M, Kreider K, Thompson J, Germanwala S, Greifenkamp J. Improving outcomes in patients with peripheral arterial disease. J Vasc Nurs. 2018;36:166–172. doi: 10.1016/j.jvn.2018.06.005

4. Swaminathan A, Vemulapalli S, Patel MR, Jones WS. Lower extremity amputation in peripheral artery disease: improving patient outcomes. Vasc Health Risk Manag. 2014;10:417–424. doi: 10.2147/VHRM.S50588

5. Sustained effect of intensive treatment of type 1 diabetes mellitus on development and progression of diabetic nephropathy: the Epidemiology of Diabetes Interventions and Complications (EDIC) study. Jama. 2003;290:2159–2167.

6. Iyer SR, Annex BH. Therapeutic Angiogenesis for Peripheral Artery Disease: Lessons Learned in Translational Science. JACC Basic Transl Sci. 2017;2:503–512. doi: 10.1016/j.jacbts.2017.07.012

7. Khan TA, Sellke FW, Laham RJ. Gene therapy progress and prospects: therapeutic angiogenesis for limb and myocardial ischemia. Gene therapy. 2003;10:285–291.

8. Kusumanto YH, van Weel V, Mulder NH, Smit AJ, van den Dungen JJ, Hooymans JM, Sluiter WJ, Tio RA, Quax PH, Gans RO, et al. Treatment with intramuscular vascular endothelial growth factor gene compared with placebo for patients with diabetes mellitus and critical limb ischemia: a double-blind randomized trial. Hum Gene Ther. 2006;17:683–691. doi: 10.1089/hum.2006.17.683

9. Carmeliet P. Angiogenesis in health and disease. Nat Med. 2003;9:653–660. doi: 10.1038/nm0603-653

10. Hendriks WJAJ, Elson A, Harroch S, Pulido R, Stoker A, den Hertog J. Protein tyrosine phosphatases in health and disease. FEBS Journal. 2013;280:708–730. doi: 10.1111/febs.12000

11. Mercier C, Rousseau M, Geraldes P. Growth Factor Deregulation and Emerging Role of Phosphatases in Diabetic Peripheral Artery Disease. Front Cardiovasc Med. 2020;7:619612. doi: 10.3389/fcvm.2020.619612

12. Tonks NK. Protein tyrosine phosphatases: from genes, to function, to disease. Nature reviews Molecular cell biology. 2006;7:833–846. doi: 10.1038/nrm2039

13. Wu C, Sun M, Liu L, Zhou GW. The function of the protein tyrosine phosphatase SHP-1 in cancer. Gene. 2003;306:1–12. doi: S0378111903004001 [pii]

14. Corti F, Simons M. Modulation of VEGF receptor 2 signaling by protein phosphatases. Pharmacol Res. 2017;115:107–123. doi: 10.1016/j.phrs.2016.11.022

15. Geraldes P, Hiraoka-Yamamoto J, Matsumoto M, Clermont A, Leitges M, Marette A, Aiello LP, Kern TS, King GL. Activation of PKC-delta and SHP-1 by hyperglycemia causes vascular cell apoptosis and diabetic retinopathy. Nat Med. 2009;15:1298–1306.

16. Mercier C, Brazeau T, Lamoureux J, Boisvert E, Robillard S, Breton V, Pare M, Guay A, Lizotte F, Despatis MA, Geraldes P. Diabetes Impaired Ischemia-Induced PDGF (Platelet-Derived Growth Factor) Signaling Actions and Vessel Formation Through the Activation of Scr Homology 2-Containing Phosphatase-1. Arterioscler Thromb Vasc Biol. 2021;41:2469–2482. doi: 10.1161/ATVBAHA.121.316638

17. Lizotte F, Pare M, Denhez B, Leitges M, Guay A, Geraldes P. PKCdelta Impaired Vessel Formation and Angiogenic Factor Expression in Diabetic Ischemic Limbs. Diabetes. 2013;62:2948–2957. doi: 10.2337/db12-1432

18. Croteau L, Mercier C, Fafard-Couture E, Nadeau A, Robillard S, Breton V, Guay A, Lizotte F, Despatis MA, Geraldes P. Endothelial deletion of PKCdelta prevents VEGF inhibition and restores blood flow reperfusion in diabetic ischemic limb. Diab Vasc Dis Res. 2021;18:1479164121999033. doi: 10.1177/1479164121999033

19. Paquin-Veillette J, Lizotte F, Robillard S, Beland R, Breton MA, Guay A, Despatis MA, Geraldes P. Deletion of AT2 Receptor Prevents SHP-1-Induced VEGF Inhibition and Improves Blood Flow Reperfusion in Diabetic Ischemic Hindlimb. Arterioscler Thromb Vasc Biol. 2017;37:2291–2300. doi: 10.1161/ATVBAHA.117.309977

20. Campisi J. Aging, cellular senescence, and cancer. Annu Rev Physiol. 2013;75:685-705. doi: 10.1146/annurev-physiol-030212-183653

21. Spinelli R, Baboota RK, Gogg S, Beguinot F, Bluher M, Nerstedt A, Smith U. Increased cell senescence in human metabolic disorders. J Clin Invest. 2023;133. doi: 10.1172/JCI169922

22. Herranz N, Gil J. Mechanisms and functions of cellular senescence. J Clin Invest. 2018;128:1238–1246. doi: 10.1172/JCI95148

23. Regina C, Panatta E, Candi E, Melino G, Amelio I, Balistreri CR, Annicchiarico-Petruzzelli M, Di Daniele N, Ruvolo G. Vascular ageing and endothelial cell senescence: Molecular mechanisms of physiology and diseases. Mech Ageing Dev. 2016;159:14–21. doi: 10.1016/j.mad.2016.05.003

24. Shimizu I, Minamino T. Cellular Senescence in Arterial Diseases. J Lipid Atheroscler. 2020;9:79–91. doi: 10.12997/jla.2020.9.1.79

25. Bitar MS. Diabetes Impairs Angiogenesis and Induces Endothelial Cell Senescence by Up-Regulating Thrombospondin-CD47-Dependent Signaling. Int J Mol Sci. 2019;20. doi: 10.3390/ijms20030673

26. Schneider CA, Rasband WS, Eliceiri KW. NIH Image to ImageJ: 25 years of image analysis. Nat Methods. 2012;9:671–675. doi: 10.1038/nmeth.2089

27. Robillard S, Tran K, Lachance MS, Brazeau T, Boisvert E, Lizotte F, Auger-Messier M, Boudreault PL, Marsault E, Geraldes P. Apelin prevents diabetes-induced poor collateral vessel formation and blood flow reperfusion in ischemic limb. Front Cardiovasc Med. 2023;10:1191891. doi: 10.3389/fcvm.2023.1191891

28. Lizotte F, Denhez B, Guay A, Gevry N, Cote AM, Geraldes P. Persistent Insulin Resistance in Podocytes Caused by Epigenetic Changes of SHP-1 in Diabetes. Diabetes. 2016;65:3705–3717. doi: 10.2337/db16-0254

29. Hazarika S, Dokun AO, Li Y, Popel AS, Kontos CD, Annex BH. Impaired angiogenesis after hindlimb ischemia in type 2 diabetes mellitus: differential regulation of vascular endothelial growth factor receptor 1 and soluble vascular endothelial growth factor receptor 1. Circulation research. 2007;101:948–956. doi: 10.1161/CIRCRESAHA.107.160630

30. Li HH, Cai X, Shouse GP, Piluso LG, Liu X. A specific PP2A regulatory subunit, B56gamma, mediates DNA damage-induced dephosphorylation of p53 at Thr55. The EMBO journal. 2007;26:402–411. doi: 10.1038/sj.emboj.7601519

31. Reid MA, Wang WI, Rosales KR, Welliver MX, Pan M, Kong M. The B55alpha subunit of PP2A drives a p53-dependent metabolic adaptation to glutamine deprivation. Mol Cell. 2013;50:200–211. doi: 10.1016/j.molcel.2013.02.008

32. Tsao CW, Aday AW, Almarzooq ZI, Anderson CAM, Arora P, Avery CL, Baker-Smith CM, Beaton AZ, Boehme AK, Buxton AE, et al. Heart Disease and Stroke Statistics-2023 Update: A Report From the American Heart Association. Circulation. 2023;147:e93–e621. doi: 10.1161/CIR.0000000000001123

33. Dua A, Gologorsky R, Savage D, Rens N, Gandhi N, Brooke B, Corriere M, Jackson E, Aalami O. National assessment of availability, awareness, and utilization of supervised exercise therapy for peripheral artery disease patients with intermittent claudication. J Vasc Surg. 2020;71:1702–1707. doi: 10.1016/j.jvs.2019.08.238

34. Rivard A, Silver M, Chen D, Kearney M, Magner M, Annex B, Peters K, Isner JM. Rescue of diabetes-related impairment of angiogenesis by intramuscular gene therapy with adeno-VEGF. The American journal of pathology. 1999;154:355–363.

35. Schonborn M, Laczak P, Pasieka P, Borys S, Plotek A, Maga P. Pro- and Anti-Angiogenic Factors: Their Relevance in Diabetic Foot Syndrome-A Review. Angiology. 2022;73:299–311. doi: 10.1177/00033197211042684

36. Koch S, Tugues S, Li X, Gualandi L, Claesson-Welsh L. Signal transduction by vascular endothelial growth factor receptors. Biochem J. 2011;437:169–183. doi: 10.1042/BJ20110301

37. Bhattacharya R, Kwon J, Wang E, Mukherjee P, Mukhopadhyay D. Src homology 2 (SH2) domain containing protein tyrosine phosphatase-1 (SHP-1) dephosphorylates VEGF Receptor-2 and attenuates endothelial DNA synthesis, but not migration*. Journal of molecular signaling. 2008;3:8. doi: 10.1186/1750-2187-3-8

38. Sosinska-Zawierucha P, Mackowiak B, Staniszewski R, Suminska-Jasinska K, Maj M, Krasinski Z, Breborowicz A. Sulodexide Slows Down the Senescence of Aortic Endothelial Cells Exposed to Serum from Patients with Peripheral Artery Diseases. Cell Physiol Biochem. 2018;45:2225–2232. doi: 10.1159/000488167

39. Wu H, Wu J, Zhou S, Huang W, Li Y, Zhang H, Wang J, Jia Y. SRT2104 attenuates diabetes-induced aortic endothelial dysfunction via inhibition of P53. J Endocrinol. 2018;237:1–14. doi: 10.1530/JOE-17-0672

40. Gogiraju R, Xu X, Bochenek ML, Steinbrecher JH, Lehnart SE, Wenzel P, Kessel M, Zeisberg EM, Dobbelstein M, Schafer K. Endothelial p53 deletion improves angiogenesis and prevents cardiac fibrosis and heart failure induced by pressure overload in mice. J Am Heart Assoc. 2015;4. doi: 10.1161/JAHA.115.001770

41. Sun Z, Pan X, Zou Z, Ding Q, Wu G, Peng G. Increased SHP-1 expression results in radioresistance, inhibition of cellular senescence, and cell cycle redistribution in nasopharyngeal carcinoma cells. Radiat Oncol. 2015;10:152. doi: 10.1186/s13014-015-0445-1

42. Yokoi T, Fukuo K, Yasuda O, Hotta M, Miyazaki J, Takemura Y, Kawamoto H, Ichijo H, Ogihara T. Apoptosis signal-regulating kinase 1 mediates cellular senescence induced by high glucose in endothelial cells. Diabetes. 2006;55:1660–1665. doi: 10.2337/db05-1607

43. Tonelli C, Chio IIC, Tuveson DA. Transcriptional Regulation by Nrf2. Antioxid Redox Signal. 2018;29:1727–1745. doi: 10.1089/ars.2017.7342

44. Guo H, Chen J, Yu H, Dong L, Yu R, Li Q, Song J, Chen H, Zhang H, Pu J, Wang W. Activation of Nrf2/ARE pathway by Anisodamine (654-2) for Inhibition of cellular aging and alleviation of Radiation-Induced lung injury. Int Immunopharmacol. 2023;124:110864. doi: 10.1016/j.intimp.2023.110864

45. Bao XY, Deng LH, Huang ZJ, Daror AS, Wang ZH, Jin WJ, Zhuang Z, Tong Q, Zheng GQ, Wang Y. Buyang Huanwu Decoction Enhances Revascularization via Akt/GSK3beta/NRF2 Pathway in Diabetic Hindlimb Ischemia. Oxid Med Cell Longev. 2021;2021:1470829. doi: 10.1155/2021/1470829

46. Faraonio R, Vergara P, Di Marzo D, Pierantoni MG, Napolitano M, Russo T, Cimino F. p53 suppresses the Nrf2-dependent transcription of antioxidant response genes. J Biol Chem. 2006;281:39776–39784. doi: 10.1074/jbc.M605707200

47. Villeneuve NF, Sun Z, Chen W, Zhang DD. Nrf2 and p21 regulate the fine balance between life and death by controlling ROS levels. Cell Cycle. 2009;8:3255–3256. doi: 10.4161/cc.8.20.9565

48. Seshacharyulu P, Pandey P, Datta K, Batra SK. Phosphatase: PP2A structural importance, regulation and its aberrant expression in cancer. Cancer Lett. 2013;335:9–18. doi: 10.1016/j.canlet.2013.02.036

49. Du Y, Kowluru A, Kern TS. PP2A contributes to endothelial death in high glucose: inhibition by benfotiamine. American journal of physiology Regulatory, integrative and comparative physiology. 2010;299:R1610–1617. doi: 10.1152/ajpregu.00676.2009

